# *Aspergillus fumigatus* In-Host HOG pathway mutation for Cystic Fibrosis Lung Microenvironment Persistence

**DOI:** 10.1101/2021.02.17.431756

**Authors:** Brandon S. Ross, Lotus A. Lofgren, Alix Ashare, Jason E. Stajich, Robert A. Cramer

**Affiliations:** Department of Microbiology and Immunology, Geisel School of Medicine at Dartmouth, Hanover, NH 03755 USA; Department of Microbiology and Plant Pathology and Institute for Integrative Genome Biology University of California Riverside, Riverside, CA 92507 USA; Department of Medicine, Dartmouth-Hitchcock Medical Center, Lebanon, NH 03766 USA

**Keywords:** Aspergillus fumigatus, Cystic Fibrosis, chronic infection, hypoxia, osmotic stress, oxidative stress, genomics

## Abstract

The prevalence of *Aspergillus fumigatus* colonization in individuals with Cystic Fibrosis (CF) and subsequent fungal persistence in the lung is increasingly recognized. However, there is no consensus for clinical management of *A. fumigatus* in CF individuals, due largely to uncertainty surrounding *A. fumigatus* CF pathogenesis and virulence mechanisms. To address this gap in knowledge, a longitudinal series of *A. fumigatus* isolates from an individual with CF were collected over 4.5 years. Isolate genotypes were defined with whole genome sequencing that revealed both transitory and persistent *A. fumigatus* in the lung. Persistent lineage isolates grew most readily in a low oxygen culture environment and conidia were more sensitive to oxidative stress inducing conditions compared to non-persistent isolates. Closely related persistent isolates harbor a unique allele of the high osmolarity glycerol (HOG) pathway mitogen activated protein kinase kinase, Pbs2 (*pbs2^C2^*). Data suggest this novel *pbs2^C2^* allele arose *in vivo* and is necessary for the fungal response to osmotic stress in a low oxygen environment through hyperactivation of the HOG (SakA) signaling pathway. Hyperactivation of the HOG pathway through *pbs2^C2^* comes at the cost of decreased conidia stress resistance in the presence of atmospheric oxygen levels. These novel findings shed light on pathoadaptive mechanisms of *A. fumigatus* in CF, lay the foundation for identifying persistent *A. fumigatus* isolates that may require antifungal therapy, and highlight considerations for successful culture of persistent fungal CF isolates.

**Importance:** *Aspergillus fumigatus* infection causes a spectrum of clinical manifestations. For individuals with Cystic Fibrosis (CF), Allergic Bronchopulmonary Aspergillosis (ABPA) is an established complication, but there is a growing appreciation for *A. fumigatus* airway persistence in CF disease progression. There currently is little consensus for clinical management of *A. fumigatus* long-term culture positivity in CF. A better understanding of *A. fumigatus* pathogenesis mechanisms in CF is expected to yield insights into when antifungal therapies are warranted. Here, a 4.5-year longitudinal collection of *A. fumigatus* isolates identified a persistent lineage that harbors a unique allele of the Pbs2 MAPKK necessary for unique CF-relevant stress phenotypes. Importantly for *A. fumigatus* CF patient diagnostics, this allele provides increased CF lung fitness at a cost of reduced *in vitro* growth in standard laboratory conditions. These data illustrate a molecular mechanism for *A. fumigatus* CF lung persistence with implications for diagnostics and antifungal therapy.

## Introduction

The prevalence of fungal colonization in Cystic Fibrosis (CF) individual’s lungs has become increasingly recognized in recent years (1–3). CF arises from defects in the human Cystic Fibrosis Transmembrane Conductance Regulator (hCFTR) that result in a thickened mucus layer over lung epithelial cells (4). The dense layer of mucins, oligosaccharides, and extracellular DNA blocks normal mucociliary clearance, an important innate defense against environmental debris and inhaled microbes (5). A reduction in mucociliary clearance in combination with mucus’s nutrient content renders CF airways susceptible to both transient microbial colonization and persistent infection (6). Although decidedly different in duration and evolutionary potential, both of these host-microbe interactions lead to development of recurrent inflammation, steep oxygen and osmolarity gradients, and progressive fibrotic tissue damage resulting in lung function decline (7–10).

It is well-documented that several fungal genera that cause severe infections in immune compromised patient populations are also commonly isolated from CF patient airways (11–13). *Aspergillus fumigatus* has been identified as the most commonly isolated filamentous fungal species from CF patients (1, 14). Estimates point to *A. fumigatus* colonization being present in up to ∼60% of CF patients, and recent data demonstrates that initial colonization occurs as early as infancy (15, 16). *Aspergillus*- specific diseases affecting CF patients include *Aspergillus* Bronchitis (AB) and Allergic Bronchopulmonary Aspergillosis (ABPA), each of which are associated with persistent fungal colonization and worse patient outcomes (17). Despite evidence that *A. fumigatus* persistence can last for years in CF patients, and reports that persistent fungal colonization and associated AB and ABPA negatively impact patient outcomes, there is little consensus for the importance of direct clinical management of *Aspergillus* culture positivity in CF patients (18–20). This is in contrast to bacterial persistence, where dynamic and persistent bacterial populations are widely understood to contribute to worsening outcomes in CF (21, 22). Consequently, CF patients are routinely treated with antibiotics, but far less frequently with antifungals even in the face of persistent fungal colonization (23). A better understanding of the pathogenesis and virulence mechanisms of *A. fumigatus* in CF patients is expected to help identify in what contexts antifungal therapies should be considered.

Analyses of CF patient bacterial isolates has been key to understanding CF bacterial pathogenesis and virulence in CF disease progression. For example, genomic and genetic analyses of CF bacterial isolates revealed significant intraspecies diversity, phenotypic convergence, and in-host evolution in microbes such as *Pseudomonas aeruginosa* and *Burkholderia* species among others (24, 25). While intraspecies diversity in *A. fumigatus* is associated with a spectrum of disease manifestations, there is little consensus as to whether *A. fumigatus* culture positivity in CF patients is a sign of infection or transient colonization (18, 26). Until recently, efforts to examine longitudinal genotypic diversity in CF *A. fumigatus* isolates have relied on lower-resolution genotyping techniques in contrast to bacterial whole-genome sequence analyses (27–29). Moreover, phenotypic analysis of *A. fumigatus* CF isolates has largely focused on antifungal drug susceptibility and resistance (30). A recent study from Engel and colleagues reported parasexual recombination as a potential mechanism for persistence of *A. fumigatus* in chronic infections, but the implications of this genetic diversity for phenotypes outside of antifungal resistance are unclear (31). There remains a lack of longitudinal studies that directly address the genotypic and CF-relevant phenotypic characteristics of persistent *A. fumigatus* isolates. Longitudinal studies have significant potential to shed light on CF-specific *A. fumigatus* pathogenesis mechanisms similar to studies in CF associated bacteria (32).

Here, we defined the genotypes and phenotypes of a longitudinal series of *A. fumigatus* isolates collected from a CF patient over the course of ∼4.5 years (the AF100 series). Whole-genome sequencing and phylogenetic analyses of 29 isolates from 12 different time points revealed extensive genetic diversity and two lineages of persistent isolates. One lineage of 5 persistent isolates has evidence of positive natural selection. Isolates from this persistent lineage revealed conidia with a striking increased sensitivity to growth in ambient oxygen conditions (normoxia) and oxidative stress-inducing culture conditions, but also an increase in osmotic stress resistance compared to non-persistent isolates. Conidial phenotypes result in part from a unique allele of *pbs2*, the MAP kinase kinase associated with the high osmolarity glycerol (HOG) MAPK pathway that most likely arose *in vivo* during growth in the CF airway (33, 34). Further, we observe that this unique *pbs2* allele promotes hyphal osmotic and oxidative stress resistance at the cost of conidial oxidative stress susceptibility. Taken together, these results reveal new insights into pathoadaptive mechanisms of persistent *A. fumigatus* CF isolates, and provides support for the importance of further genetic and phenotypic studies of *A. fumigatus* CF isolates in the context of disease progression.

## Materials and Methods

### Strain isolation and Media

All growth assays were performed using a modified *Aspergillus* Minimal Medium (AMM) as described previously by Cove (35). Briefly, 10g D-glucose (1% w/v) was dissolved in 20ml of 50x Salt Solution, 1 ml Trace Elements Solution, 66.666 ml .3M L-glutamine solution (as nitrogen source, 20 mM final), and ddH_2_O to 1L (pH adjusted to 6.5 using NaOH). All media were made using 1.5% (w/v) agar (Beckton-Dickinson, Franklin Lakes, NJ) and autoclaved for at least 20 mins at 121 °C. 20 ml plates were poured after addition of filter sterilized FeSO_4_ solution (70 mM, 2 mM final concentration). Standard growth conditions were 72 hours at 37 C, 21% O_2_, and 5% CO_2_ in the dark, and all plate images were taken using a Canon Powershot SX40 HS camera, using automatic settings. Morphological characteristics were assessed as previously described (36). The liquid minimal medium used for mycelial DNA cultures consisted of 1% (w/v) D-Glucose, 0.5% Yeast Extract (w/v, Beckton-Dickinson), 20 ml 50x Salt Solution, 1 ml Trace Elements solution, and NaNO_3_ to 20 mM. Liquid media was sterilized as described above.

All CF clinical isolates used in this study were cultured from expectorated sputum, collected from the AF100 patient during hospital visits (Fig 1A). Sputum samples were collected and stored at -80 °C by the Clinical Microbiology Laboratory at Dartmouth-Hitchcock Medical Center (DHMC) in collaboration with the Translational Research Core at Dartmouth. Isolates were obtained from the AF100 samples in one of two ways: 1) as individual colonies sub-cultured onto Sabouraud Dextrose agar (Sab) slants following primary sputum culture on either Sab or blood agar by staff at the Clinical Microbiology Laboratory at DHMC, or 2) as a collection of multiple colonies picked from sputum cultured on Sab plates for 30hrs in the Cramer laboratory. All isolates were grown in pure culture for 3 days at 37 °C, followed by single spore isolation, propagation, and storage in -80C glycerol stocks (25%). Single spores were collected by cutting individual germinated spores after 16 hours of growth on AMM plates at 30 °C and transferred to individual plates for further culture. All AF100 isolates were initially confirmed to be *A. fumigatus* using Sanger sequencing of PCR products from the Internal Transcribed Spacer (ITS1) region according to Schoch, following spore DNA extraction as described by Fraczek (37, 38).

**Figure 1:**
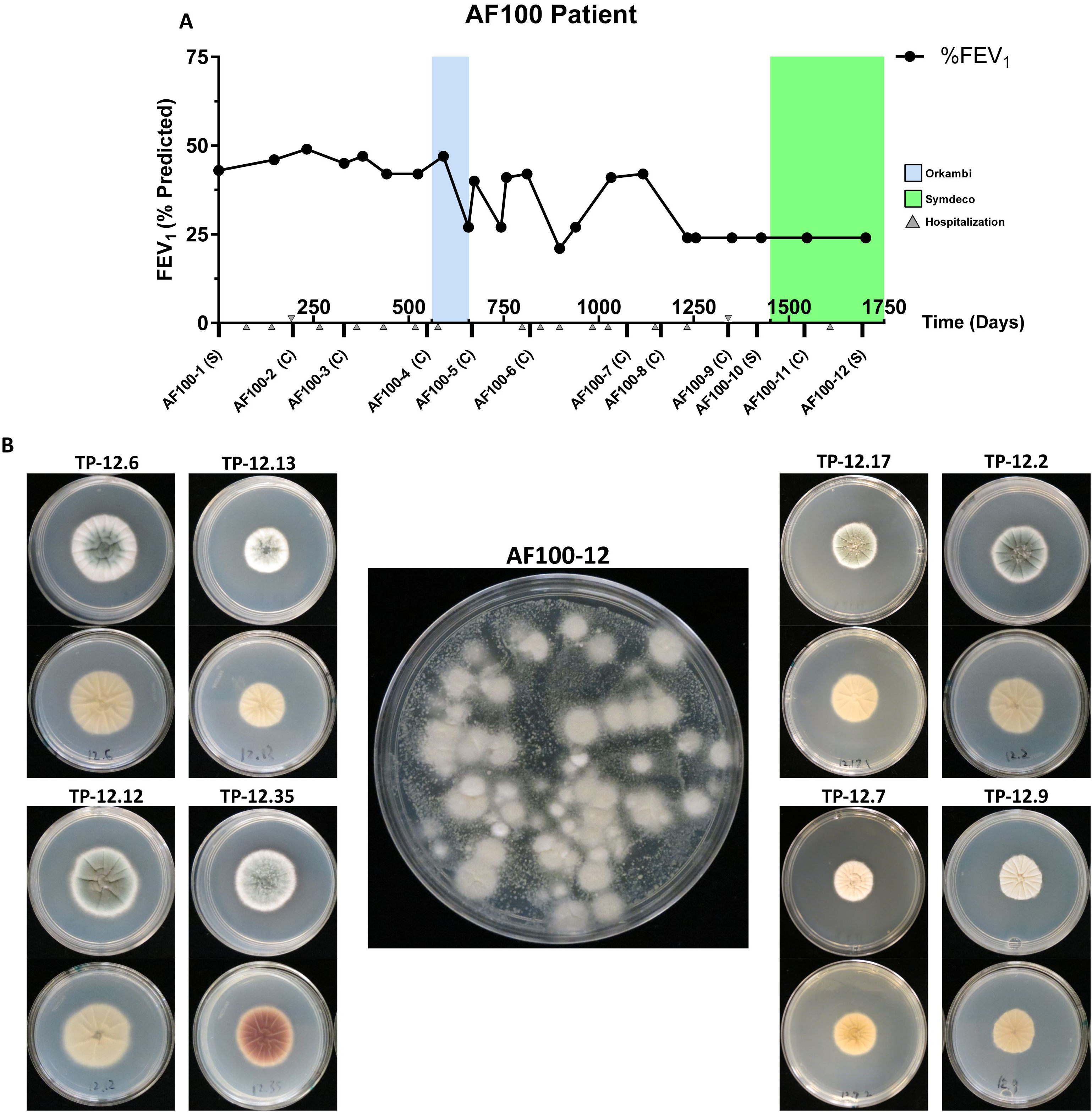
The longitudinal *Aspergillus fumigatus* isolate series from the AF100 cystic fibrosis individual reveals morphological diversity. A) Timeline of sample isolation, Forced Expiratory Volume in 1 second (FEV_1_), and cystic fibrosis medication history for the AF100 individual over the time covered in this study. The letter below each AF100 sample represents the sample type: C = individual/sub cultured colony sample from clinical microbiology laboratory at Dartmouth Hitchcock Medical Center, Sp = sputum sample. B) Single colony images highlighting morphology of front (top image) and back (bottom image) of *Aspergillus* minimal medium (AMM) plates for representative colonies picked from plated AF100-12 sputum , inset. Sputum was initially cultured on Sabouraud Dextrose Agar for 30-hours in normoxia, and single cultures were subsequently cultured on AMM for 48 hours at 37° C in normoxia.

### Genome sequencing and assembly

Genomic DNA (gDNA) was extracted from 24hr mycelia grown in station petri plates using liquid media described above, according to previously published methods (39). DNA concentration was quantified using a Qubit 2.0 Fluorometer (Invitrogen) and the manufacturer’s recommended Broad Range protocol. Genomic sequencing was caried out on either Illumina NovaSeq 6000 or NextSeq 500 machines (Table S1). DNA sequencing libraries were prepared using either the NEBNext Ultra II DNA Library Prep Kit (for NovaSeq 6000 sequenced genomes), or the SeqOnce DNA library kit utilizing Covaris mechanized shearing (for the NextSeq 500 sequenced isolates), both following manufacturer recommendations with paired end library construction and barcoding for multiplexing.

**Table 1.** Genes and mutations of interest from Clade 2 isolates.

| Gene ID | Product Description (FungiDB) | Type of Mutation | Nucleotide Change |
| --- | --- | --- | --- |
| **Afu1g07160**<br>*(ubp3)* | Ortholog(s) have ubiquitin-specific protease activity, role in protein deubiquitination and cytosol, nucleus localization | Splice donor variant | 522+2T>G |
| **Afu1g15950**<br>*(pbs2)* | **Putative mitogen-activated protein kinase (MAPKK)** | **Missense variant** | **590C>T** |
| **Afu2g10620** | Ortholog(s) have protein serine/threonine kinase activity, protein serine/threonine kinase inhibitor activity | Missense variant | 199G>A |
| **Afu3g05900**<br>**(*ste7*)** | MAPK kinase (MAPKK) | Start site lost | 2T>C |
| **Afu3g08010**<br>**(*ace1*)** | C2H2 transcription factor | Missense variant | 2175G>C |
| **Afu3g10080** | Ortholog(s) have cytosol localization | Missense variant | 531C>G |
| **Afu3g11940** | Has domain(s) with predicted zinc ion binding activity | Stop codon gained | 1711C>T |
| **Afu3g11970**<br>**(*pacC*)** | C2H2 finger domain transcription factor | Frameshift variant | 1628dupC |

### Phylogeny of strains

Aligned SNPs from all isolates (identified as described below) were used to construct a phylogeny employing the Maximum Likelihood algorithm IQ-TREE (40). The best fit model for evolutionary rates was determined to be GTR+F+ASC according to the BIC score assessed with the ModelFinder function in IQ-TREE. The phylogeny was inferred under this model and 1000 rapid bootstrap iterations. Individual branch support values were assessed using a Shimodaira-Hasegawa approximate likelihood ratio test (SH-aLRT) with 1000 iterations. Tree visualization and trait mapping was carried out using the program Iroki (41).

### Phenotypic Assays using CF-relevant stresses

Phenotypic profiling assays were performed with cultures growing on AMM media supplemented as indicated with treatments that induce CF-relevant stresses. The conditions tested included ambient oxygen (normoxia, 21% O_2_, 5% CO_2,_ 74% N_2_), low oxygen (hypoxic, 1% O_2_, 5% CO_2_, 94% N_2_ ), osmotic (2M sorbitol), oxidative (25 μM menadione), and azole antifungal drug (0.2 μg/ml voriconazole) stresses (42, 43). For conidial-based assays, conidia were collected from 3-day old cultures, counted using a hemocytometer, and diluted to 500 spores/μl using 0.01% Tween 80. 2 μl drops were inoculated onto plates and grown for 72 hours under the indicated conditions. For hyphal-based assays, conidia were plated as above and grown for 48 hours in normoxia, except for TP-12.7 which was grown in hypoxia. A sterile .5 cm cork-borer was used to transfer hyphal plugs from the leading, non-sporulating, edge of the colonies to new plates with or without the indicated stresses and plates were grown for 72 hours in normoxia or hypoxia. Colony diameter was measured by averaging 2 perpendicular diameter measurements per plate. To compare the susceptibility of the isolates to a given stress, we quantified the absolute value of the change in colony diameter compared to non-stressed growth conditions, or Δ^CD^ ( = | [mean of untreated] – [mean of treated replicate] |, averaged across all replicates). Assays were performed in biological triplicate and statistical analysis was performed in PRISM using one-way ANOVA with a multiple comparison post-test.

### Comparative Genomics

To examine the genetic diversity of the AF100 isolates in the context of global *A. fumigatus* diversity, raw sequence reads from 68 strains of *A. fumigatus* were downloaded from the NCBI database. These strains were combined with the 53 strains sequenced as part of this project, including 29 AF100 isolates (total number of strains analyzed = 121, Table S1). The sequence reads for each strain were aligned to the AF293 reference genome downloaded from FungiDB v.46 (44, 45) using BWA v0.7.17 (46) and converted to the BAM file format using samtools v1.10 (47). Duplicate reads were marked for ignoring and the BAM files indexed using picard tools v2.18.3 (http://broadinstitute.github.io/picard). To avoid overcalling variants near alignment gaps, reads were realigned using RealignerTargetCreator and IndelRealigner in the Genome Analysis Toolkit GATK v3.7 (48). Variants (SNPs and INDELS) were genotyped relative to AF293 using HaplotypeCaller in GATK v4.0 (49). Filtering was accomplished using GATK’s SelectVariants with the following parameters: for SNPS: -window-size = 10, - QualByDept < 2.0, -MapQual < 40.0, -QScore < 100, -MapQualityRankSum < -12.5, - StrandOddsRatio > 3.0, -FisherStrandBias > 60.0, -ReadPosRankSum < -8.0. For INDELS: -window-size = 10, -QualByDepth< 2.0, -MapQualityRankSum < -12.5, - StrandOddsRatio > 4.0, -FisherStrandBias > 200.0, -ReadPosRank < -20.0 , - InbreedingCoeff < -0.8. Resultant variants were annotated with snpEff (50) relative to the AF293. Variants that overlapped transposable elements (TEs) were removed by positional mapping to locations of annotated TEs in the FungiDB v.46 release of AF293 using BEDtools -subtract (51). For all analyses except the mutational accumulation analysis, variants annotated in snpEff as “intergenic”, “intron_variant”, and “synonymous_variant” were excluded from the analysis. Mutational accumulation analysis necessarily included variants annotated as “synonymous_variant”.

Variants were analyzed using a custom script implemented in the R programing environment (52) and assessed in three ways (52). First, to identify variants relevant to the tested phenotypes observed in the clade isolates, we targeted clade variants relative to four strains that failed to demonstrate these phenotypes under culture assay: the transient clinical strains TP-2, TP-7, and TP-11.1, isolated as part of this study, along with the non-clinical strain IF1SW-F4 previously isolated from the International Space Station, hereafter referred to as the “out group” isolates (53). Second, to identify variants exclusive to each clade, we targeted variants in clade isolates relative to all non-clade isolates in the dataset (n = 109). Finally, to look for recent signatures of adaptation to the human lung environment, variants in the Clade 1 isolates were assessed relative to the putatively ancestral isolate TP-1.3, which was isolated at the first time point and fell directly outside of Clade 1 in the phylogenomic analysis.

### Positive Selection Analysis

Using the outgroup targeted variant set, mutation accumulation curves were constructed for each clade using a custom R script (AF100_variant_analysis.R) that analyzed only on those variants that had gone to fixation (present in a given time point, and all subsequent time points for that clade). Significance was assessed using linear regression at P < 0.05 in the R programing environment.

### Construct design, generation, and transformation for allelic exchange

To generate the *pbs2* allele swap mutants, we designed primers (Table S2) and used fusion PCR to generate a construct using the *pbs2* genomic sequence from either TP-12.7 (*pbs2^C1^*) or TP-9 (*pbs2^C2^* Fig 4A). We used the Prime Star high-fidelity Taq polymerase and buffer (Takara Bio USA, Mountain View, CA) with 50ng of template DNA and the following thermocycler settings: 95 °C for 3mins, followed by 35 cycles of 95 °C for 10 s, 55 °C for 15 s, and 72 °C for 1 min/kb of amplicon, finishing with 72 °C for 10min. Constructs were confirmed via gel electrophoresis and extracted using the GeneJET Gel Extraction Kit (ThermoFisher Scientific, Waltham, MA). We used these constructs for transformation via homologous recombination in TP-9 protoplasts generated with lysing enzymes from *Trichoderma* spp. (Sigma Aldrich, St. Louis, MO, USA). Transformation was performed according to previously published protocols (36).

### Protein extraction and Western blotting

Mycelia from 30-hour stationary liquid AMM cultures (1×10^6^ spores/ml) were used for media transfer and RNA extraction. Mycelia were transferred to either fresh AMM or AMM with 1M NaCl for 15 minutes and then flash frozen, lyophilized overnight, and bead beaten for 1 min with 2.3 mm beads. Protein was extracted using urea-based extraction buffer (per 10ml: 9M Urea, 500ul 20% SDS, 25mM Tris HCl pH7.5, 1mM EDTA, 100μl 10% Igepal, 10μl 1M DTT, 100μl HALT Protease inhibitor cocktail [ThermoScientific], 100mM PMSF, ddH_2_O to 10ml). 1ml of buffer was added to powdered mycelia, vortexed, and tubes were centrifuged at 13,000rpm for 5 minutes. ∼500 μl were removed, diluted with 200 μl of buffer, and protein quantified via Bradford analysis. 40 μg of protein was used for each sample for Western blot analysis. Proteins were transferred from a 10% SDS-PAGE gel onto a nitrocellulose membrane for a Western blot assay using the Trans-Blot turbo transfer system (Bio-Rad, Hercules CA). Phosphorylated SakA (P- SakA) was detected using a rabbit anti-P-p38 antibody #9211 (Cell Signaling Technology, Danvers MA) at 1:1,000 dilution. Fluorescent imaging and quantification were performed using the LI-COR Odyssey CLx system and manufacturer’s protocol, along with REVERT total protein stain for normalization (LI-COR Biosciences, Lincoln NE).

### RNA extraction and qRT-PCR

Mycelia from 30-hour stationary liquid AMM cultures (1×10^6^ spores/ml) were used for media transfer and RNA extraction. Mycelia were transferred to either fresh AMM or AMM with 1M NaCl for 15 minutes and then flash frozen, lyophilized overnight, and bead beaten for 1 min with 2.3 mm beads. Homogenate was suspended in 1 ml of Trisure (Meridian Biosciences, Cincinnati OH) and RNA was extracted as previously described (54). For qRT-PCR, 5 μg of RNA was treated with the Ambion Turbo DNAse kit (ThermoFisher Scientific, Waltham MA) according to the manufacturer’s protocol, and DNase-treated RNA was reverse transcribed as previously described (54). Statistical analysis was performed with one-way ANOVA with Dunnet post-test for multiple comparisons. Data were collected on a CFX Connect Real-Time PCR Detection System (Bio-Rad, Hercules CA) with CFX Maestro Software.

### Data Availability

All genome data generated as part of this project was deposited into the NCBI Short Read Archive under BioProject no. PRJNA666940. The code and datafiles for variant assessment associated with this project can be found on GitHub in the repository: https://github.com/stajichlab/AF100 (DOI: 10.5281/zenodo.4409345).

## Results

### The AF100 series is a longitudinal series of Aspergillus fumigatus isolates from a single CF patient

We investigated a series of *A. fumigatus* isolates collected from a person living with a F508del/F508del CFTR genotype, herein called the AF100 series. The AF100 series is an ongoing longitudinal collection of >30 isolates collected from patient sputum over a ∼4.5-year period (Fig 1A). Over the course of the study, the patient exhibited a steady decrease in %FEV_1_ and multiple hospitalizations associated with persistent *A. fumigatus* culture positivity but was not treated with antifungal therapy. The patient began consistent *A. fumigatus* culture positivity starting in the year 2005. Sputum samples for each of the 12 timepoints are designated as “AF100-X”, where X corresponds to the timepoint in the series. Fungal isolates were obtained from the AF100 sputum samples in one of two ways: as individual colonies sub-cultured from primary sputum cultures by staff at the Clinical Microbiology Laboratory at Dartmouth-Hitchcock Medical Center (DHMC), and as a collection of multiple different colonies picked from patient sputum samples cultured in the Cramer laboratory (labeled C and S respectively, Fig. 1A). Isolates from the AF100 samples were subsequently designated as “TP-X”, so that TP-2 is the isolate from the AF100-2 sample, and TP-3 is from AF100-3, etc. It is important to note sputum samples that allowed us to pick multiple isolates (AF100-1, AF100-10, AF100-11, and AF100-12) required further numbering to differentiate isolates (i.e. TP-12.7, TP-12.9, TP-12.13, etc.).

We were initially curious to understand the level of *A. fumigatus* diversity in an individual sputum sample; to do this, we used a preliminary qualitative screen for morphological diversity in the AF100-12 sputum sample. After culturing the sputum on Sabouraud’s Dextrose agar medium for 30 hours and sub-culturing individual colonies to *Aspergillus* Minimal Medium (AMM) for 48 hours, we observed a high degree of morphological diversity (Fig 1B). We recovered at least 8 unique colony morphotypes from the AF100-12 sputum sample, with striking variation in coloration, furrowing, aerial hyphae, and colony diameter (Fig 1B). These observations are consistent with previous reports on the presence of multiple phenotypically diverse *A. fumigatus* isolates in the CF lung at any given time (28, 55, 56). To better understand these observations and determine potential genetic relationships amongst the AF100 isolates we turned to whole genome sequencing.

### Whole-Genome Sequencing and phylogenetic analyses reveal significant genetic diversity and 2 persistent lineages within the AF100 series

Genomic DNA from 29 AF100 series isolates (6 from AF100-1, 1 each from AF100-2 through AF100-9, 2 from AF100-10, 3 from AF100-11, and 10 from AF100-12) was sequenced with Illumina based technology. Genomes were sequenced to an average depth of 55x coverage (Table S1). To better understand the genetic relationships between the AF100 series isolates and generate a phylogeny, we used the 29 genomes from the AF100 series and a collection of *A. fumigatus* genomes from non-CF clinical and environmental isolates (n = 92) including 68 publicly available genomes and 53 newly sequenced genomes. Across the 121 strains we identified 223,369 variants relative to the AF293 reference genome (77,644 excluding intergenic, synonymous, and intronic variants). Of the total variants, 85,304 represent Single Nucleotide Polymorphisms (SNPs), which were used to construct a 121-strain *A. fumigatus* ML phylogeny that yielded a well-supported topology with most branches possessing > 90% bootstrap support (Fig. 2A). AF100 isolates occupy ∼15 distinct positions across the tree, indicating the AF100 series represents a genetically diverse population.

**Figure 2:**
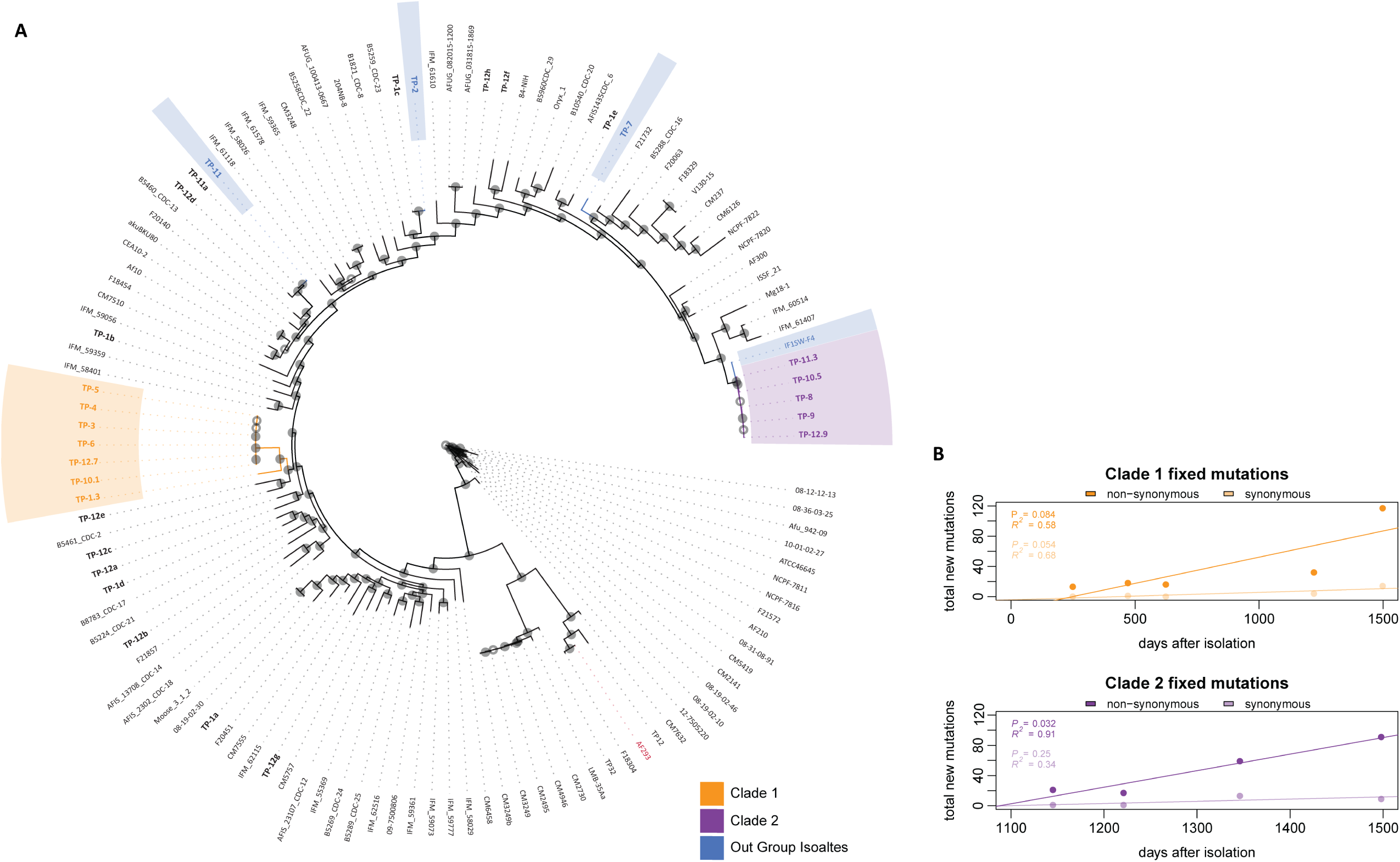
Whole-genome single nucleotide polymorphism phylogeny of 121 *Aspergillus fumigatus* isolates confirms genetic diversity and reveals 2 persistent clades in the AF100 series. A) Illumina Whole-Genome sequencing and phylogenomic analyses generated a well-supported topology using 121 *A. fumigatus* isolates (29 from AF100). AF100 isolates are designated as “TP-X.Y”, where X indicates the sample number and Y identifies different isolates from the same sample (where applicable). Clade 1 isolates are highlighted in orange, and Clade 2 isolates in purple. Isolates from outside these Clades used for outgroup analysis are highlighted in blue, the AF293 reference is highlighted in red, and all AF100 isolates are listed in bold. Filled circles at bipartition’s represent > 90% bootstrap support, open circles represent < 90%. The data supports a history of multiple colonization events for the AF100 patient, including both transient and persistent isolates. B) Mutation accumulation curves for fixed synonymous and nonsynonymous mutations in the Clade 1 and Clade 2 isolates. Only those mutations acquired and then found in all subsequent isolates were considered in each Clade. Significance was assessed using linear regression at P < 0.05 in the R programing environment. Clade 2 showed a significant accumulation of nonsynonymous mutations relative to synonymous mutations, suggesting Clade 2 is under positive selection.

While many isolates from the AF100 series occupy their own positions in the tree and do group into a phylogenetic clade, 11 of the isolates cluster into two distinct clades, indicating repeated isolation of similar genotypes from the AF100 patient over the 4.5- year time course (Fig 2A). We designated these CF patient-specific clades as Clades 1 and 2. Both Clades consist of isolates from timepoints that span large periods of time, with 6 isolates in Clade 1 ranging from AF100-3 to AF100-12 (∼3.5 years) and 5 isolates in Clade 2 ranging from AF100-8 to AF100-12 (∼1.5 years) (Fig 2A, Fig S1). The close genetic relationship between the isolates within each Clade, as well as the timescale between isolations, supports the possibility that each Clade represents repeated isolation of a persistent genotype from the AF100 individual. Further, both Clades feature isolates from the AF100-10 and AF100-12 plated sputum samples, which suggests they are present at the same time in the AF100 individual (Fig. 1A, 2A).

Although AF100 isolates that fell outside of Clades 1 and 2 were generally dispersed across the phylogeny, a single isolate from the AF100-1 sputum sample is most closely related to isolates of Clade 1 (Fig 2A). Isolation of TP-1.3 pre-dates the first Clade 1 isolate by ∼1 year (Fig. 1A) and considering the comparatively few variants that distinguish these two isolates (∼1757 mutations, Table S1) TP-1.3 may represent an ancestral isolate to Clade 1. Although no clear ancestor was identified for Clade 2 within the AF100 series, the environmental isolate IF1SW-F4 isolated from the International Space Station (53) surprisingly differs by only 643 total mutations (Fig 2A, Table S1). These results indicate that the AF100 series represents a strikingly diverse population of *A. fumigatus* isolates, featuring at least 2 distinct genotypes that are either persistent or recurrent colonizers.

To further investigate the likelihood of these Clades representing persistent genotypes compared to recurrent colonizers, we performed mutation accumulation analysis using the variant data from the different isolates of each Clade. Analysis compared the number of fixed, nonsynonymous variants in each Clade over time to the number of fixed, synonymous variants in each Clade over time. A significant accumulation of nonsynonymous variants supports a given Clade’s genotype as potentially being under positive selection, in this case potentially facilitating persistence in the AF100 patient (57). Our analysis revealed a significant accumulation of nonsynonymous mutations in Clade 2 but not in Clade 1 (Fig 2B). We consequently focused on Clade 2 isolates to interrogate CF-relevant phenotypic characteristics of this *A. fumigatus* lineage.

### Clade 2 conidia have unique phenotypic profiles in response to CF-relevant stresses

In the presence of representative CF lung microenvironment conditions, Clade 2 isolates had unique and adapted phenotypes compared to the control strains TP-2 (non-clade strain) and IF1SW-F4 (closest to Clade 2 sequenced environmental strain). A striking phenotype for Clade 2 isolates is reduced growth in normoxic (21% O_2_, 5% CO_2_, 74% N_2_) compared to hypoxic (1% O_2_, 5% CO_2_, 94% N_2)_ conditions. Some Clade 2 colonies grew nearly double in size in hypoxic conditions, resulting in Δ^CD^ values ranging from 1.68 to 1.95 (Fig. 3A). Clade 2 isolates had significantly decreased growth in normoxic compared to hypoxic conditions compared to the control strains IF1SW-F4 (0.58) and TP-2 (0.32). When tested for growth in presence of the triazole antifungal voriconazole, Clade 2 isolates were significantly more resistant than outgroups as evidenced by substantial colony growth on solid AMM containing 0.2 ug/ml voriconazole (Fig 3B). Clade 2 Δ^CD^ ranged from 0.55 to 0.63, significantly lower than TP-2 (1.47) and IF1SW-F4 (1.88, Fig 3B). All isolates had voriconazole MICs lower than the clinical resistance threshold as measured by CLSI microbroth dilution (< 1 ug/ml, Fig. S2), but were capable of growth on solid medium containing sub-MIC levels of drug. The finding that Clade 2 conidia grow well at sub-MIC levels of voriconazole is particularly striking given that medical records indicate the AF100 patient has not received azole therapy since being treated at DHMC (1996, ∼24 years).

**Figure 3:**
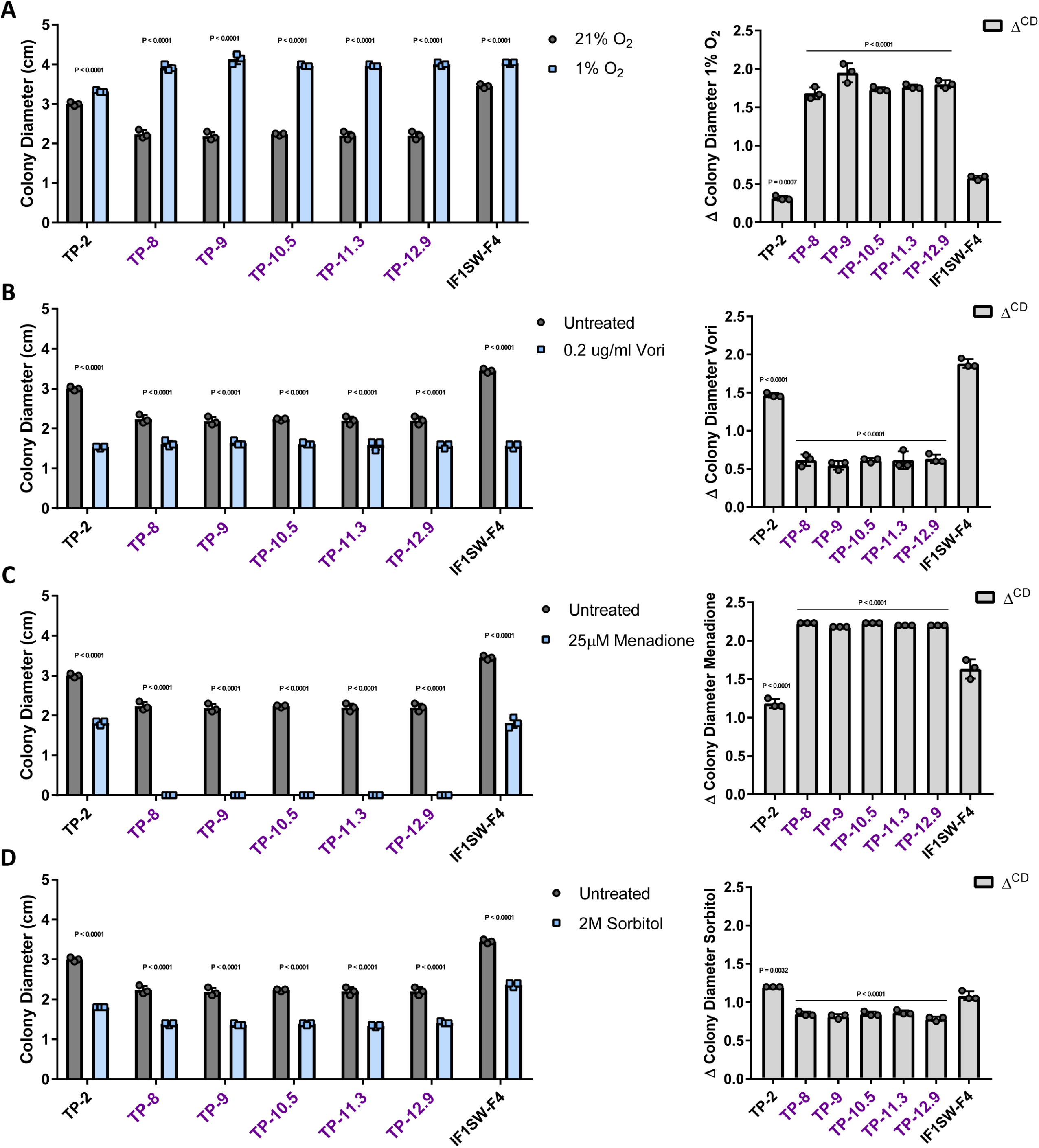
Clade 2 isolate conidia show altered CF-relevant stress phenotypes. Colony diameter was calculated by taking the average of perpendicular colony measurements, and Δ^CD^ was calculated as the absolute value of the mean of colony diameter of the untreated replicates minus colony diameter of a treated replicate and averaged across all replicates for a given strain and condition. Each was calculated after conidial inoculation and 72 hours of growth on AMM under A) hypoxic (1% O_2_) B) sub- MIC voriconazole (0.2 μg/ml) C) menadione-induced (25μM) oxidative and D) sorbitol- induced (2M) osmotic stress conditions, normalized to untreated growth. For colony diameter measurements, individual symbols represent the average colony diameter of 3 technical replicates. For Δ^CD^, symbols represent the Δ^CD^ calculated per-replicate, as assessed according to the described methods. Error bars represent standard deviation; for colony diameter, significance was assessed via 2-way ANOVA using Sidak multiple comparison post-test to compare treated vs untreated for each isolate, and for Δ^CD^ via 1- way ANOVA using Dunnett multiple comparisons post-test to compare all isolates to the IF1SW-F4 outgroup (using Prism 7, significance cutoff P<0.05). These findings indicate Clade 2 isolates have altered CF-relevant stress resistance profiles relative to outgroups.

Conidia from Clade 2 isolates were significantly more susceptible to the superoxide-generating compound menadione with Δ^CD^ ranging from 2.18 to 2.23 compared to 1.18 for TP-2 and 1.63 for IF1SW-F4 (Fig 3C). Strikingly, Clade 2 isolates were not able to germinate at the menadione concentration tested, a phenotype not shared with any other AF100 isolate (Fig 3C). Lastly, compared to TP-2 and IF1SW-F4, conidia from Clade 2 isolates were significantly more resistant to sorbitol-mediated osmotic stress, with Δ^CD^ ranging from 0.78 to 0.87 across the Clade compared to 1.2 for TP-2 and 1.08 for IF1SW-F4 (Fig 3D). Together, these results suggest that conidia from Clade 2 isolates have unique phenotypes for CF-relevant stresses relative to the controls, which suggests that Clade 2 isolates are uniquely adapted to the complex oxidative and osmotic environment found in CF airways (8).

### Comparative genomic analyses reveal mutations in stress response pathways that are unique to Clade 2

To identify alleles associated with the CF stress phenotypes observed in Clade 2 isolates, we first utilized comparative genomic analyses. Compared to the AF293 reference genome, we identified 23,218 variants present in Clade 2 isolates. Variant comparisons relative to the outgroup isolates (TP-2, TP-7, TP-11.1, and IF1SW-F4) reduced this set to 643 variants in Clade 2. Because this method allows for possible miscalls and single deletions in various isolates, we further restricted the variant set to include only those appearing in all Clade 2 isolates. This final set of 88 variants comprised 38 confident conserved single base deletions and 2 spanning deletions (Supplemental File 1). To prioritize variants of interest that may be associated with specific Clade 2 phenotypes, we further filtered the variants and identified 38 exclusive to Clade 2 and found in no other isolates across the 121-isolate phylogeny. These Clade 2 exclusive variants included 4 SNV deletions and 6 indels (Supplemental File 2). Clade 2- exclusive variants were found in 37 unique genes and one gene, Afu5g06915, with 2 variants. Interestingly, there were no overlapping variants exclusive to both Clade 1 and Clade 2 (found in both clades, but in no other isolates across the phylogeny).

GO term enrichment via the FungiDB database revealed that of the 37 genes with unique mutations in Clade 2, six are annotated with roles in stress sensing and response: Afu1g07160 (*ubp3*), Afu1g15950 (*pbs2/B*), Afu2g10620, Afu3g05900 (*ste7*), Afu3g08010 (*sltA/ace1*), and Afu3g11940 (45). Specifically, Pbs2 and Ste7 are well-studied MAPKKs involved in extracellular stress responses. Pbs2 has roles in response to hypoxic, oxidative, and osmotic stresses as part of the well-studied HOG MAPK pathway in fungi (58–60); Ste7 is the MAPKK of the mating/invasive growth (MpkB) pathway and is responsive to nutrient conditions and the presence of other fungi (61). Ubp3 and Ace1/SltA have been demonstrated to interact with Ste7 (in other fungal species) and downstream MpkB pathway members, respectively; Ace1/SltA is involved in salt stress and is responsive to voriconazole (62–65). Afu3g10080 and Afu3g11970 (*pacC*) are also mutated uniquely in Clade 2 and have annotated roles in response to intracellular stress, the latter of which is a known virulence determinant and internal pH stress-response factor and works concomitantly with Ace1/SltA (66). These 8 genes and their mutations of interest are summarized in Table 1. Given the oxidative and osmotic stress phenotypes observed in Clade 2 isolates, the cytosine to thymine point mutation at position 590 in the coding sequence of *pbs2* (*pbs2^C2^*) was investigated further.

### The pbs2^C2^ allele is necessary for CF-relevant stress phenotypes in TP-9 conidia

The 590 point mutation is one of 2 nonsynonymous SNPs in Pbs2^C2^ compared to the AF293 reference genome, and the allele results in a unique P197L amino acid substitution in a region homologous to a domain necessary for MAPK interaction in *S. cerevisiae* (Fig. 4A) (67). To test the hypothesis that *pbs2^C2^* was necessary for the observed Clade 2 phenotypes, we conducted an allelic exchange experiment using the Clade 2 isolate TP-9 and the *pbs2* allele from Clade 1 to generate strain TP-9*^pbs2C1^* (*pbs2^C1^*, Fig 4A). The TP-9*^pbs2C1^* strain was confirmed via Southern blot analysis and Sanger sequencing. As predicted, the TP-9*^pbs2C1^* mutant conidia had a significantly altered stress resistance profile relative to the TP-9 background (Fig 4B-E). TP-9*^pbs2C1^* grew significantly better in normoxic conditions as reflected by the smaller Δ^CD^ than TP-9 (∼0.93 vs ∼1.95, respectively) and also phenocopied TP-12.7, the source of the *pbs2^C1^* allele (Fig 4B). Significant conidiation was also restored in TP-9*^pbs2C1^* (Fig. 4B). Additionally, TP-9*^pbs2C1^* conidia were capable of growth on menadione in contrast to TP- 9, with an overall lower Δ^CD^ (∼1.42 vs ∼2.18 respectively, Fig 4D); this growth in the presence of menadione is even more striking considering TP-9 conidia do not germinate in these conditions. Together with the hypoxia:normoxia colony growth, these data indicate that the *pbs2^C2^* allele is necessary for the observed oxygen sensitivity of Clade 2 conidia. Considering the above change in oxygen sensitivity and the importance of oxygen in ergosterol biosynthesis, we also examined voriconazole resistance of the TP- 9*^pbs2C1^* mutant. As expected, TP-9*^pbs2C1^* showed a decrease in voriconazole resistance relative to the no drug control and TP-9 (Fig 4E).

We lastly examined the osmotic stress resistance of the TP-9*^pbs2C1^* strain conidia compared to TP-9. We expected to observe an increase in osmotic stress resistance in TP-9*^pbs2C1^* strain conidia based on the oxidative stress phenotype and known contributions of the HOG pathway to both osmotic and oxidative stress responses. However, TP-9*^pbs2C1^* conidia showed a severe growth defect in the presence of 2M sorbitol; with low growth in the treated condition, TP-9*^pbs2C1^*Δ^CD^ (∼2.75) was significantly larger than that of TP-9 and TP-12.7 as well as IF1SW-F4, the non-CF isolate closely related to Clade 2 (Fig 4F). The conidia osmotic stress susceptibility in the absence of *pbs2^C2^* suggests that the *pbs2^C2^* allele confers increased conidial osmotic stress resistance in the CF lung environment at the cost of reduced oxidative stress resistance. Taken together, the phenotypic profiling data indicates the *pbs2^C2^* allele is necessary for the unique CF-relevant conidial phenotypes found in Clade 2.

### Pbs2^C2^ is not associated with menadione susceptibility in hyphae, but is necessary for osmotic resistance independent of growth stage

The severe Clade 2 isolate oxygen sensitivity presented a conundrum if these were isolates persisting in the CF lung. How could these isolates evade the robust oxidative burst utilized by leukocytes to prevent *A. fumigatus* infection to establish and progress disease? We thus considered that hyphal stress responses may reveal more about the importance of the *pbs2^C2^* allele in *A. fumigatus* persistent CF infections. We modified the conidial stress assay to use 0.5cm hyphal plugs from established colonies as the starting inoculum. Similar to the observed normoxia:hypoxia growth phenotype with conidia, TP-9 hyphae had a significantly higher Δ^CD^ than TP-9*^pbs2C1^* (Fig 5A). This result indicates that, as with conidia, the TP-9 hyphae have reduced growth in normoxia; however, the magnitude of this phenotype is not as large as observed with conidia (Fig 4A, 5A). Surprisingly, however, TP-9 hyphae were unaffected by menadione in normoxia and in hypoxia only showed a minor reduction in growth (Fig 5B). This result with TP-9 hyphae is in striking contrast to TP-9 conidia that did not germinate at the same menadione concentration. Despite the comparable resistance of the TP-9*^Ppbs2C1^* conidia to menadione, the TP-9*^Ppbs2C1^* hyphae were slightly more susceptible than TP-9 hyphae in normoxia, and slightly more resistant in hypoxia (Fig 5B). Thus, while the *pbs2^C2^* allele does alter the response to menadione in hyphae, the effect size between TP-9 and TP-9*^pbs2C1^* suggests that *pbs2^C2^* allele is not directly necessary for superoxide mediated stress resistance in TP-9 hyphae (Fig 5B). These data support the conclusion that the *pbs2^C2^* allele is more detrimental to conidial responses to oxygen stress than in mature hyphae.

**Figure 4:**
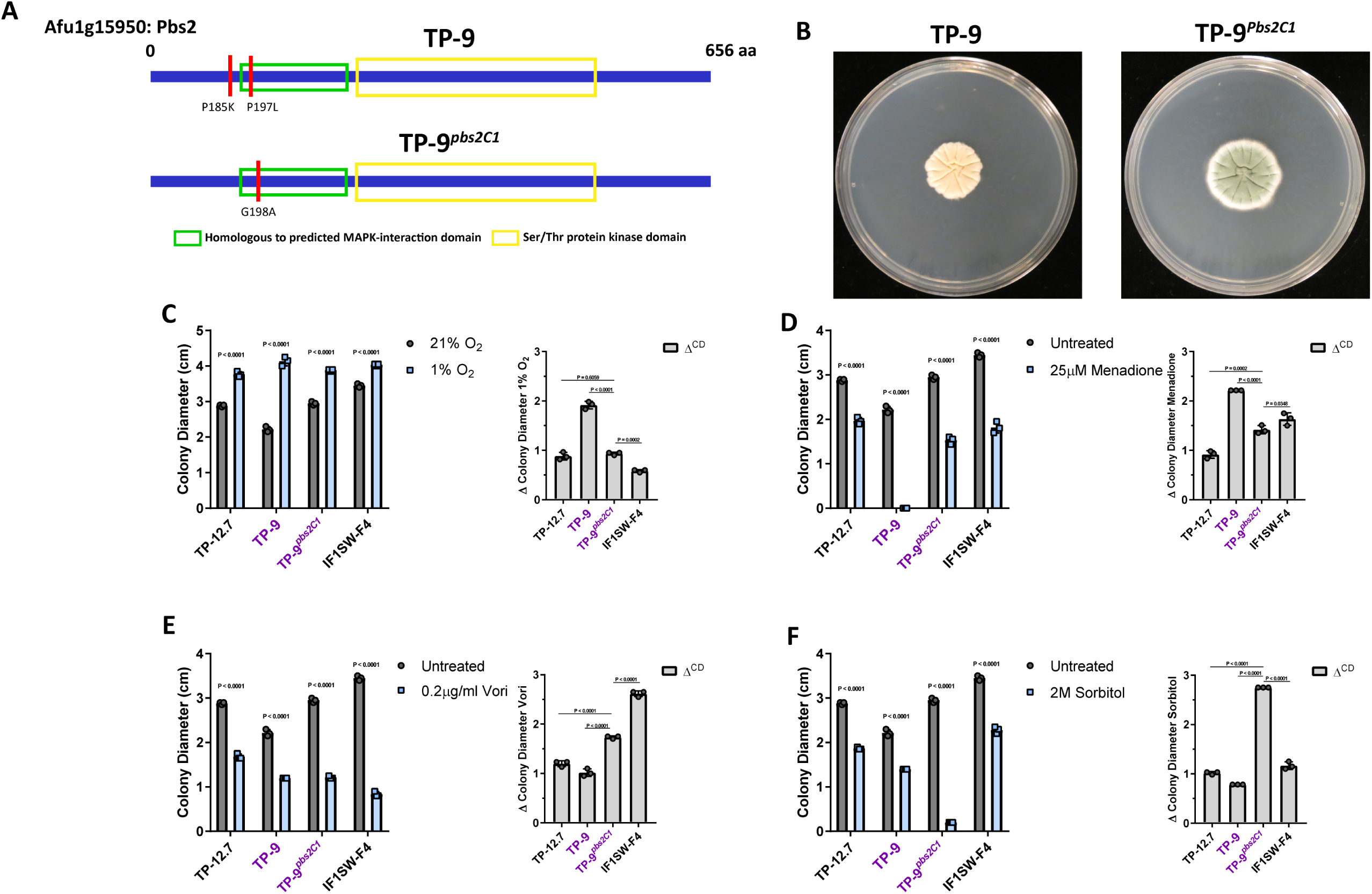
Allelic replacement of *pbs2^C2^* to generate TP-9*^pbs2C1^* results in morphological changes and significantly altered conidial stress responses. A) schematic representation of the Pbs2 protein domains in *A. fumigatus*, with variants in each of the indicated isolates marked in red (4 total nucleotide changes). B) TP-9*^pbs2C1^* displays morphological and growth changes in normal conditions (72 hours at 37 °C in normoxia). For conidial phenotyping, symbols represent the average colony diameter and Δ^CD^ as described previously. For colony diameter, significance was assessed via 2- way ANOVA using Sidak multiple comparison post-test to compare treated vs untreated for each isolate, and for Δ^CD^ via 1-way ANOVA using Dunnett multiple comparisons post- test to compare all isolates to the TP-9*^pbs2C1^* mutant (using Prism 7, significance cutoff P<0.05). TP-9*^pbs2C1^* displayed significantly altered profiles for the Clade 2 oxygen phenotypes, including partially rescued oxygen sensitivity in both hypoxia (C) and oxidative stress (D) conditions. Interestingly, TP-9*^pbs2C1^* showed a slight decrease in voriconazole resistance (E). Most notably, TP-9*^pbs2C1^* mutant showed a significant growth defect in the presence of osmotic stress (F), suggesting the *pbs2^C2^* allele may be important for persistence of Clade 2.

**Figure 5:**
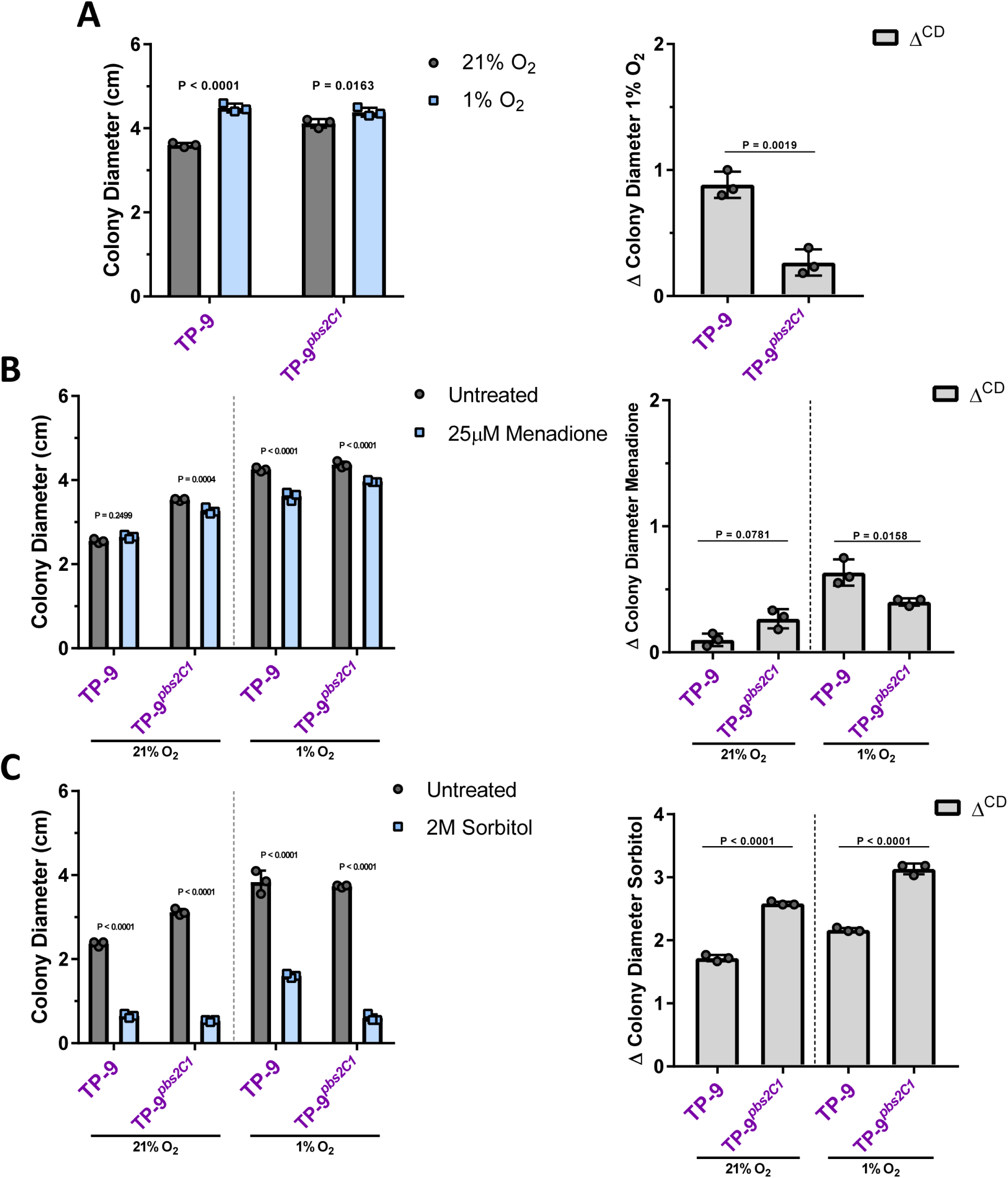
*Pbs2^C2^* is not associated with severe oxidative stress susceptibility in hyphae but is necessary for osmotic stress resistance. Hyphal phenotypic stress assays used 0.5 cm^2^ hyphal plugs cut from 48-hour cultures and grown for an additional 72 hours at 37 °C. Individual symbols represent the average colony diameter and the average Δ^CD^ as described previously. For colony diameter, significance was assessed via 2-way ANOVA using Sidak multiple comparison post-test to compare treated vs untreated for each isolate, and for Δ^CD^ via 1-way ANOVA using Tukey multiple comparisons post-test to compare all isolates based on oxygen conditions (using Prism 7, significance cutoff P<0.05). TP-9 hyphae had lower Δ^CD^ under hypoxic (A) and oxidative (B) stress conditions than conidia, but similar trends held up when compared to TP-9*^pbs2C1^* hyphae. C) Osmotic stress conditions resulted in poor growth for TP-9*^pbs2C1^* compared to TP-9, suggesting *pbs2^C2^* is necessary for resistance.

We next examined the osmotic stress sensitivity of TP-9 hyphae. Regardless of oxygen conditions, TP-9 hyphae are significantly more resistant to osmotic stress relative to TP-9*^pbs2C1^* hyphae (Fig 5C). These data suggest *pbs2^C2^* is necessary for osmotic stress resistance in TP-9 regardless of fungal growth stage. The magnitude of the osmotic stress sensitivity in TP-9*^pbs2C1^* hyphae is larger in hypoxia than normoxia, further suggesting that the *pbs2^C2^* allele is important for Clade 2 stress responses in CF-like conditions. Overall, these results strongly suggest that the *pbs2^C2^* allele results in increased hyphal stress resistance at the cost of conidial oxygen sensitivity in Clade 2.

### The pbs2^C2^ allele is not sufficient to replicate Clade 2 oxygen or osmotic stress phenotypes in Clade 1

We next asked if the *pbs2^C2^* allele is sufficient to induce the unique Clade 2 oxidative and osmotic stress phenotypes in another genetic background. We chose TP- 12.7, as it is the most recent isolate from Clade 1 and is the source of the *pbs2^C1^* allele. We performed this allelic exchange according to the same construct design using *pbs2^C2^* to generate the TP-12.7*^pbs2C2^* strain (Fig S3). We used conidia and hyphae from TP-12.7 and the TP-12.7*^pbs2C2^* strain to examine growth in hypoxic, oxidative, and osmotic stress conditions. There was no significant difference between TP-12.7 and the TP-12.7*^pbs2C2^* strain for hypoxic growth with conidia or hyphae. Interestingly, the extreme Clade 2 conidia menadione sensitivity phenotype was also not observed in TP-12.7*^pbs2C2^,* indicating that the *pbs2^C2^* allele is not sufficient to induce conidia menadione susceptibility in the Clade 1 genetic background (Fig S3). Further, there was no significant difference between the strains for osmotic stress. These data suggest genetic interactions with other allelic variants in the Clade 2 background are necessary to fully express the unique Clade 2 Pbs2-mediated stress responses.

### Pbs2^C2^ is necessary for full HOG pathway activity in response to osmotic stress in Clade 2

In order to understand the effect of the Clade 2 *pbs2* allele on HOG pathway function, we first tested our isogenic set of strains for sensitivity to the drug fludioxonil. In fungi, resistance to fludioxonil requires loss of HOG pathway activity (68, 69). We observed that TP-9 was more susceptible to fludioxonil compared to TP-9*^pbs2C1^* in both normoxic and hypoxic conditions, which suggests that TP-9 has increased HOG pathway activity relative to TP-9*^pbs2C1^* (Fig 6A). To further test if Pbs2^C2^ is required for HOG pathway activity in response to stress, we examined phosphorylation of SakA, the *A. fumigatus* HOG MapK homolog via quantitative fluorescent western blots. We observed that, relative to the TP-9*^pbs2C1^* mutant, TP-9 had significantly more P-SakA signal upon NaCl induced osmotic stress, which supports the hypothesis that *pbs2^C2^* is necessary for increased HOG pathway activity in response to osmotic stress in Clade 2 (Fig 6B). These data also suggest a molecular basis for the increased osmotic stress resistance in TP-9 relative to the related environmental isolate IF1SW-F4 (Fig 6B).

**Figure 6:**
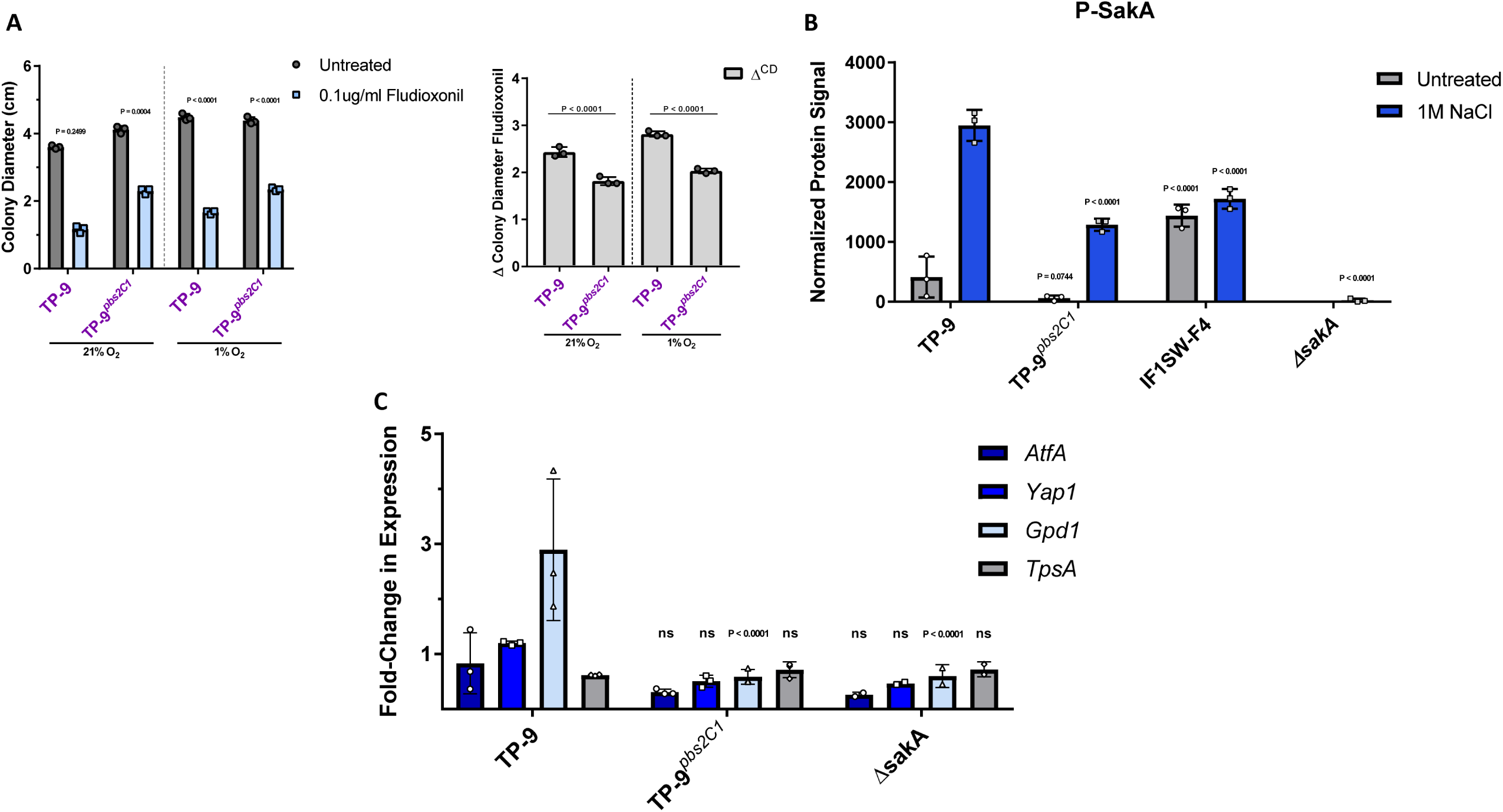
Loss of *pbs2^C2^* results in reduced HOG pathway activity in response to osmotic stress. A) Using the hyphal stress assay as previously described, TP-9*^pbs2C1^* is significantly more resistant to growth on AMM + 0.1 μg/ml fludioxonil, which suggests reduced HOG pathway activity. For colony diameter, significance was assessed via 2- way ANOVA using Sidak multiple comparison post-test to compare treated vs untreated for each isolate, and for Δ^CD^ via 1-way ANOVA using Tukey multiple comparisons post-test to compare isolates based on oxygen conditions (using Prism 7, significance cutoff P<0.05). B) Quantification from LI-COR fluorescent Western blotting for P-SakA (the active form of the Pbs2 target). Bars represent the average of 3 biological replicates (symbols), normalized to the REVERT total protein stain protocol (LI-COR Biosciences). Statistical analysis was performed using 2-way ANOVA with Dunnett multiple comparison post-test to compare all isolates to TP-9 at the same conditions, and error bars represent standard deviation. TP-9*^pbs2C1^* has lower P-SakA levels in both untreated and 1M NaCl conditions, which suggests that the *pbs2^C2^* allele is involved in altering HOG pathway activity. C) mRNA levels of HOG pathway associated target genes. *pbs2^C2^* qRT-PCR was performed using primers for the actin (*actA*) and β-tubulin (*tubA* housekeeping genes as well as a selection of downstream SakA (HOG) targets (*atfA, yap1, gpd1,* and *tpsA*). RNA extracted from mycelia in mid-log phase transferred to fresh AMM or AMM + 1M NaCl for 15 minutes. Bars represent the average of 3 biological replicates (symbols) normalized to the fresh AMM group. Statistical analysis was performed using 2-way ANOVA with Dunnett multiple comparison post-test to compare all isolates to TP-9 on a transcript-by-transcript basis, significance cutoff P<0.05 (ns indicates P>0.13). The lower response by TP-9*^pbs2C1^* suggests it has reduced SakA signaling in response to osmotic stress.

To further test our hypothesis that *pbs2^C2^* is necessary for increased HOG pathway activity in Clade 2, we examined expression of 4 putative SakA (HOG) target genes in response to NaCl osmotic stress (70). The HOG target genes were chosen based on their roles in response to different stresses: Afu3g11330 (*atfA*) a bZIP transcription factor required for oxidative and heat stress resistance, Afu6g09930 (*yap1*) a key regulator of the oxidative stress response, Afu2g08250, a homolog of *gpd1* necessary for glycerol biosynthesis and osmotic stress resistance in *S. cerevisiae*, and Afu6g12950 (*tpsA*) that is required for trehalose biosynthesis in response to temperature stress (71–74).

TP-9 hyphae were able to mount a strong transcriptional response to osmotic shock, as indicated by increased mRNA levels of *gpd1* (Fig 6C). The specificity and necessity of this response for osmotic stress resistance is shared across a number fungal species (75). However, loss of the *pbs2^C2^* allele resulted in significantly reduced *gpd1* induction, and no change in other examined HOG pathway target gene mRNA levels. This loss of *gpd1* mRNA levels was similar to levels observed in the Δ*sakA* control strain, further suggesting TP-9*^pbs2C1^* has reduced HOG pathway activity compared to the CF isolate TP-9 (Fig 6C). Taken together, the unique *pbs2^C2^* allele is necessary for Clade 2 conidial oxidative stress susceptibility and growth stage-independent osmotic stress resistance via increased SakA HOG pathway activity.

## Discussion

*Aspergillus fumigatus* colonization in Cystic Fibrosis (CF) is a growing concern as recent studies and better diagnostic approaches indicate a greater degree of prevalence than previously appreciated (16, 76). An outstanding question is how *A. fumigatus* persistent presence in the CF lung contributes to disease progression. Previous studies of CF bacterial isolates identified substantial genetic diversity and specific mutations that drive pathogenesis in the CF airways (reviewed in: Folkesson et al., 2012; Winstanley et al., 2016); these studies have greatly increased our knowledge of CF bacterial pathogenesis mechanisms and rationale for empirical antibiotic therapies. In contrast, far less is known about the genotypic and phenotypic diversity of CF *A. fumigatus* isolates. A greater understanding of *A. fumigatus* colonization, persistence, and pathogenesis mechanisms is expected to help clarify when and if to apply antifungal therapies in CF patients. Here, we observed both transitory and persistent *A. fumigatus* isolates from sputum of a CF patient collected longitudinally over 4.5 years. Phenotypic analyses of the persistent isolates revealed increased fitness in response to osmotic stress and decreased fitness in ambient oxygen concentrations compared to the closely related environmental isolate IF1SW-F4. Genotypic and genetic analyses discovered an important role for the HOG MAP Kinase pathway via novel mutations in the MAPKK *pbs2,* which drove the emergence of these stress response phenotypes. These data support hypotheses that CF patients can be persistently infected with specific *A. fumigatus* genotypes and that the fungus acquires adaptive mutations to promote fitness in the dynamic CF lung environment (78, 79).

Our genomic analyses of the AF100 patient *A. fumigatus* isolates are consistent with previous studies of *A. fumigatus* intraspecies diversity. Specifically, CF patients are apparently colonized by more than one *A. fumigatus* genotype over time (80). Genomic analysis of the *A. fumigatus* isolates from the AF100 patient revealed extensive genetic diversity among the 29 isolates that occupy ∼15 distinct positions in our phylogeny of ∼121 diverse isolates (Fig 2). Genetic differences between individual isolates in our study range from a few hundred to tens of thousands of variants, with a total of 223,369 variants relative to the reference AF293 genome featured across the phylogeny. These data lend support to the long-standing hypothesis that numerous *A. fumigatus* genotypes are capable of colonizing the CF lung (55, 80). However, an important consideration for our and previous studies is that we cannot rule out that genotypes which appeared one time in our longitudinal sampling are only transitory colonizers or environmental contaminants. These data further emphasize the importance of repeated isolation of specific, related genotypes in informing diagnosis of a potential infection (and thus treatment) versus transitory colonization or environmental contamination.

Two genotypically distinct clades of persistent isolates were identified in the current study. Intriguingly, evidence for positive selection was only observed in the Clade 2 isolates. We cannot rule out that Clade 1 isolates previously experienced a positive selection event prior to our sampling or are newer to the CF patient lung environment. Further studies of Clade 1 isolates are needed to better understand their genotypic and phenotypic profiles, but it is also possible that Clade 1 and 2 isolates occupy distinct microenvironments in the CF lung. A significant accumulation of fixed non-synonymous mutations in Clade 2 over time suggests selective pressure from persistence in the CF lung environment (57). Moreover, compared to the closely-related environmental isolate IF1SW-F4, Clade 2 isolates had significantly increased fitness to CF-relevant osmotic stress and a preference for growth in a low oxygen atmosphere in addition to increased antifungal drug resistance (8, 81). These phenotypes are consistent with a genotype that has been shaped in part by long term persistence in the CF lung environment.

The high osmolarity glycerol (HOG) mitogen activated protein kinase (MAPK) pathway is known to mediate fungal cellular responses to hypoxic, oxidative, antifungal, and osmotic stresses among others (82–85). Moreover, HOG pathway mutation in clinical fungal isolates has been previously documented and associated with altered stress responses (86, 87). Consequently, the presence of the unique *pbs2^C2^* allele found only in Clade 2 isolates in this study was chosen to interrogate its role in the observed Clade 2 isolate phenotypes. Through allelic exchange experiments, we observed that *pbs2^C2^* is necessary for HOG pathway signaling in response to osmotic stress in both conidia and hyphae of Clade 2 isolates. Given the relatively ubiquitous nature of osmotic pressure in CF mucus plugs it follows that osmotic stress acts as a strong selective pressure in CF *A. fumigatus* initial colonization and long-term persistence (88).

Additional evidence that the *pbs2^C2^* allele arose *in vivo* in response to the CF lung environment is suggested by the conidia- and hyphal- specific phenotypes of Clade 2 isolates. For example, Clade 2 conidia were extremely susceptible to menadione induced oxidative stress and this was dependent on the *pbs2^C2^* allele (Fig 4). In contrast, the lack of severe oxidative stress sensitivity in Clade 2 hyphae is likely more relevant than the conidial phenotype in the context of CF persistence. A number of studies have reported visualization of hyphae and hyphal fragments in expectorated sputum from CF patients, which suggests certain isolates grow and develop in the CF lung environment (89, 90). Further, conidial production is an energetically costly process and typically does not occur *in vivo* unless in a pulmonary cavity (91). Lastly, although aerosol transmission of *A. fumigatus* between CF patients has been reported, it is unknown whether this is predominantly aerosolized hyphal fragments or conidia (29). Taken together, these data suggest the enhanced osmotic stress resistance of Clade 2 isolates comes at the expense of conidial fitness and thus likely arose *in vivo*.

Clade 2 isolates also displayed a significant growth reduction under ambient atmospheric oxygen conditions that was also *pbs2^C2^* dependent. This observation has potential clinical ramifications when it comes to culture of sputum, or bronchial lavage, samples from CF patients. Given the reduced growth rate of these CF lung adapted compared to the non-persistent isolates, and the clinical microbiology practice of growing fungal cultures under atmospheric oxygen conditions, one can speculate that persistent *A. fumigatus* isolates could be missed in clinical samples. Moreover, it was clear from our sputum sampling that picking a single *A. fumigatus* colony from a sputum plate may not reflect the true genotypic and phenotypic potential of isolates in a given patient’s airways. Further studies are needed to further test this hypothesis that has significant ramifications for detecting *A. fumigatus* infections in CF patients and in other suspected manifestations of aspergillosis (92).

Molecular details on how the specific P197L mutation in Pbs2^C2^ promotes increased fitness in the CF lung environment remain to be determined. However, the P197L mutation occurs in a region homologous to a predicted MAPK interaction domain in *S. cerevisiae* (67). Although this region of Pbs2 has not been shown to be involved in upstream or downstream HOG MAPK signaling in *A. fumigatus*, our data with fludioxonil strongly suggests the former. Fludioxonil antifungal activity in *A. fumigatus* is dependent on the Type-III Hybrid Histidine Kinase (HHK) TcsC (69). One of a number of putative histidine kinases that can activate the HOG pathway in *A. fumigatus,* TcsC signals specifically through the Ypd1-Ssk2/22-Ssk1 arm of the HOG pathway (58). Because loss of *pbs2^C2^* results in fludioxonil resistance in Clade 2 isolates, Pbs2^C2^ likely promotes HOG pathway activation through the TcsC-Ypd1-Ssk2/22/1 arm. However, we cannot rule out unique genetic interactions with other observed signaling mutations in the Clade 2 strains. In support of this alternative hypothesis, the *pbs2^C2^* allele was not sufficient to promote increased osmotic stress resistance or conidial oxidative stress sensitivity in the Clade 1 genetic background (Fig. S4). Thus, additional genetic interactions involving the *pbs2^C2^* allele in the Clade 2 background await further investigations. For example, the variants associated with *ste7* in these isolates is an intriguing observation for further study.

A primary rationale for this study was informing best practices around if and when to use antifungal therapy in CF patients positive for *A. fumigatus* cultures (23, 93). Although recent studies have observed positive effects with voriconazole therapy in ABPA patients, in CF patients antifungal treatment varies widely across patients and centers (94). Drug interactions often preclude prolonged use of both azoles and CF medications, and the indeterminate clinical importance of treatment of *A. fumigatus* in CF further complicates treatment decisions (95). Moreover, studies have isolated azole-resistant isolates from CF patients, often following initial azole treatment (96). In this regard, although the HOG pathway has been shown to be involved in antifungal resistance, the voriconazole resistance phenotype in Clade 2 is surprising given that the AF100 patient has not been treated with azoles (97). Previous studies in CF patients have also identified azole resistant *A. fumigatus* isolates from azole naïve patients (98). While the *pbs2^C2^* allele did not significantly change the microbroth dilution MIC of Clade 2 isolates, it does contribute to increased hyphal voriconazole resistance as evidenced by increased fungal growth in the presence of MIC level voriconazole (Fig 4). The observation of azole resistance and tolerance without prior azole therapy has also been observed in *Clavispora (Candida) lusitaniae* CF patient isolates (99). Thus, *in vivo* adaptation to the CF lung environment per se may promote increased antifungal drug tolerance across fungi associated with the CF lung, warrants further mechanistic study, and further emphasizes the importance of identifying persistent fungal isolates in CF patients. Taken together, our results argue that further understanding of *A. fumigatus* pathoadaptive mechanisms in CF is crucial for helping answer long standing clinical questions regarding *A. fumigatus* and cystic fibrosis.

## Acknowledgments

This work was supported by the efforts of R.A.C through funding by a pilot award from the Cystic Fibrosis Foundation (CFF) Research Development Program (STANTO15RO), a CFF basic research award (CRAMERGO19), and the NIH National Institute of Allergy and Infectious Diseases (NIAID) (grant nos. R01AI130128 and R01AI146121). A.A. is supported by NIH/NHLBI award R01HL122372. J.E.S. is a CIFAR fellow in the program Fungal Kingdom: Threats and Opportunities. Computational analyses were performed on the University of California-Riverside HPCC supported by grants from the National Science Foundation (MRI-1429826) and NIH (S10OD016290). BR was supported in part by an NIH/NHLBI T32 HL134598-01 at Dartmouth (O’Toole, GA). Core facility support provided by NIH grant P30-DK117469, NIH grant P20- GM113132 (Dartmouth BioMT COBRE), the CFF Research Development Program, and NCI Cancer Center Support Grant 5P30CA023108-37. We thank Dr. Deborah Hogan, co-director of the Dartmouth Translational Research Core, and all the staff at the Dartmouth Hitchcock Medical Center (DHMC) clinical microbiology laboratory for facilitating access to clinical samples over the course of the study.

---

**Supplemental Figure 1:** Single colony images and normoxia growth measures for the isolates from Clades 1 and 2. 1×10^3^ conidia from the 7 isolates in Clade 1 (A) or the 5 isolates in Clade 2 (B) were grown on AMM plates for 72hrs at 37°C in normoxia. C) Colony diameter measurements were taken at 72hrs and significance was assessed via 1-way ANOVA with Dunnett multiple comparisons posttest, comparing to either TP-1.3 or TP-8 for Clade 1 and 2, respectively.

**Supplemental Figure 2:** Minimum Inhibitory Concentration (MIC) of voriconazole for a select group of isolates from this study. Standard microbroth dilution using voriconazole diluted in LAMM media in 96 well plates were used, along with 1×10^5^ conidia per well from the indicated isolates. Plates were incubated for 24 hours and MIC was determined as the concentration of the first well with no visible fungal growth. 2 biological replicates are shown.

**Supplemental Figure 3:** Conidial (A-C) and hyphal (D-F) phenotyping of the TP- 12.7*^pbs2C2^* mutant reveals *pbs2^C2^* is not sufficient to reproduce the Clade 2 phenotypes. Using conidia, TP-12.7*^pbs2C2^* showed no significant difference in Δ^CD^ under either hypoxia (A) or osmotic stress conditions (C) and was actually more resistant to menadione (B) compared to TP-12.7. Using hyphae, there was no significant difference for any stress (D-F), indicating *pbs2^C2^* is not sufficient to replicate Clade 2 oxygen or osmotic stress phenotypes. Assays and statistical analyses were performed as described in Fig 5.

**Supplemental Table 1.** Whole Genome sequencing information on isolates in this study

**Supplemental Table 2.** Primer sequences used in this study.

Supplemental File 1. Text file with list of variants found in all Clade 2 isolates.

Supplemental File 2. Text file of Clade 2 exclusive variants

